# 2D Effects Enhance Precision of Gradient-Based Tissue Patterning

**DOI:** 10.1101/2023.03.13.532369

**Authors:** Yuchong Long, Roman Vetter, Dagmar Iber

**Affiliations:** Department of Biosystems Science and Engineering, ETH Zürich, Mattenstrasse 26, 4058 Basel, Switzerland; Swiss Institute of Bioinformatics, Mattenstrasse 26, 4058 Basel, Switzerland

**Keywords:** patterning, precision, morphogen gradient, reaction-diffusion model, 2D

## Abstract

Robust embryonic development requires pattern formation with high spatial accuracy. In epithelial tissues that are patterned by morphogen gradients, the emerging patterns achieve levels of precision that have recently been explained by a simple one-dimensional reaction-diffusion model with kinetic noise. Here, we show that patterning precision is even greater if transverse diffusion effects are at play in such tissues. The positional error, a measure for spatial patterning accuracy, decreases in wider tissues but then saturates beyond a width of about ten cells. This demonstrates that the precision of gradient-based patterning in two- or higher-dimensional systems can be even greater than predicted by 1D models, and further attests to the potential of noisy morphogen gradients for high-precision tissue patterning.

## Introduction

Morphogen gradients encode positional information to guide tissue patterning during embryonic development [1]. In the French flag model of patterning, morphogen concentration levels determine tissue domains of different cell fates [2]. While development is highly reproducible, reported gradient shapes differ substantially between embryos [3–7]—a seemingly contradictory situation. How a high patterning precision is achieved in spite of considerable noise in morphogen kinetics is therefore an active field of research. The problem has been studied intensely, both theoretically and experimentally [3, 4, 8–11, 7, 12– 16], and recently gained new traction with the realization that morphogen gradients may be considerably more precise than previously measured [17–19].

Early theoretical models assumed a linear gradient shape [2, 20], but measurements of a wide range of morphogen gradients have since demonstrated that most morphogen gradients are better approximated by an exponential function,

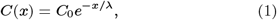

with gradient amplitude *C*_0_ and characteristic gradient length λ [4, 21, 22, 6, 7] (Fig. 1A). The position of patterning domain boundaries, herein termed the readout position,

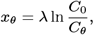

is located where the morphogen concentration reaches a threshold value *C*(*x*_*θ*_) = *C*_*θ*_. Due to molecular noise, the gradient shapes vary between embryos, and fitted exponential profiles vary in their parameters λ and *C*_0_ [4, 5, 7, 17] such that the concentration threshold is reached at different readout positions in different embryos. On average, the readout position lies at

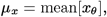

where the arithmetic mean is taken over the different embryos (or different tissue samples within an embryo). The amount of variability or dispersion in the readout positions is called the positional error. It is defined as their standard deviation [3, 4, 17]:

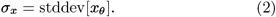

On the experimental side, most precision measurements have been carried out for the Bicoid (Bcd) gradient in the early *Drosophila* embryo [4], the Decapentaplegic (Dpp) gradient in the *Drosophila* wing disc [21, 5, 22], and Sonic Hedghehog (SHH) in the mouse neural tube [6, 7, 23]. Strikingly, in all patterning systems studied so far, the readout has been reported to be more precise than the gradients [4, 5, 7]. As a result, there has been a quest for precision-enhancing mechanisms. Spatio-temporal averaging of the morphogen concentration [4], simultaneous readout of opposing gradients [7], cell sorting [24], intermittent signaling in the *Drosophila* notum [25], and triggered self-organization in the Bcd gradient [26] have been suggested to increase gradient and readout precision. At least in the mouse neural tube, the positional error of the morphogen gradients has been overestimated [7, 17]. When corrected, the gradients are sufficiently precise to pattern the mouse neural tube. But how are such high levels of gradient precision achieved in spite of inevitable molecular noise?

**Fig. 1:**
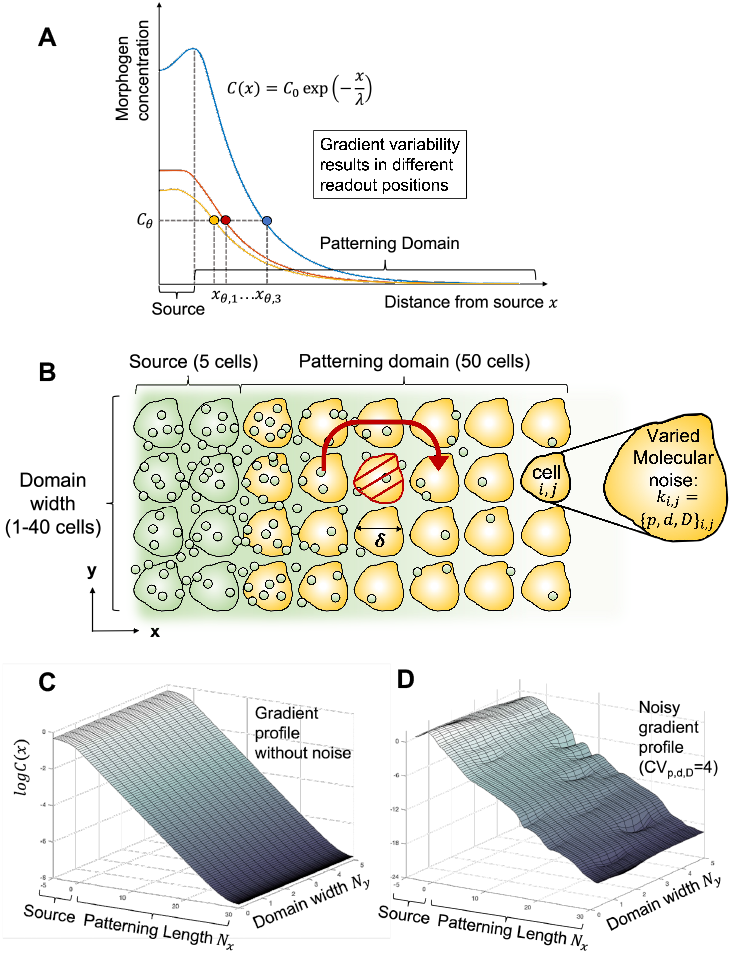
Molecular noise induces variability in gradient shape and readout positions. **A** Inter-cellular and inter-embryonic variability in the morphogen production, decay and diffusion rates leads to differently shaped gradients (colors), which attain a concentration threshold *C*_*θ*_ at different readout positions *x*_*θ*,1_…*x*_*θ*,3_. Their standard deviation is termed the positional error. **B** Morphogen diffusion in a two-dimensional tissue. Morphogens (small circles) are produced in the source (green), then secreted to the patterning domain (yellow). In 2D, morphogens can diffuse in both *x* and *y* directions, allowing them to bypass malfunctioning cells (red). In 1D, such cells have a greater impact on the gradient shape, effectively acting as transport barriers that let the morphogen concentration drop to zero beyond. **C**,**D** Simulated morphogen gradients on a 2D domain, without (C) and with molecular noise (D). The morphogen concentration *C*(*x, y*) is plotted on a logarithmic scale, and a kinetic cell-to-cell variability of CV_*p,d,D*_ = 4 (coefficient of variation) is used for illustration purposes. Cells with poor transport capabilities result in local dips in the gradient profile.

In order to analyze how molecular noise translates into gradient variability, and how the various mechanisms described above would impact precision, we previously developed a numerical framework to generate and statistically evaluate noisy morphogen gradients. An exponential gradient shape can arise from a simple 1D reaction-diffusion model with diffusion coefficient *D*, production at rate *p* in the source, and linear degradation at rate *d*:

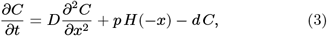

where *H*(*x*) is the Heaviside step function, limiting morphogen secretion to the source domain *x* < 0 (Fig. 1A). As a result of molecular noise, the kinetic parameters *p, d, D* will vary from cell to cell. For the reported physiological amount of parameter variability, the resulting 1D gradients differ between separate realizations (i.e., between embryos) (Fig. 1A), and have a positional error similar to that observed for the readouts in the mouse neural tube [17]. The previous analysis of 1D gradients further revealed that a small average cell diameter along the patterning axis is of key importance [18], potentially explaining why epithelia that are patterned by gradients, such as the neuroepithelium, are pseudostratified [27].

We wondered whether there would be additional effects at play that further reduce the variability of morphogen gradients. Spatio-temporal averaging over the cell surface has only minor effects on the precision of the gradient readout [18]. As a consequence, cilia and cell surface receptors can be expected to achieve a similar readout precision. Self-enhanced decay has been suggested to increase gradient robustness to noise in the morphogen source [28], but this effect is negligible when considering the effects of molecular noise more explicitly [19]. One aspect, however, has not been analyzed systematically: That of tissue dimensionality. After all, tissues are three-dimensional, and patterning domains are at least two-dimensional, even if morphogen sensing is restricted to the apical surface, as seems to be the case for SHH in the neural tube as its receptor PTCH1 is restricted to a cilium located on the apical surface [29]. One-dimensional domain approximations are known to overestimate the variability of gradients as morphogens can bypass defective cells in 2D and 3D [5] (Fig. 1B). How this effect translates into the spatial accuracy of patterning, and how it depends on the width of the patterning domain has, however, not been studied.

By extending our stochastic gradient analysis to 2D tissues (Fig. 1B–D), we now find that the readout precision and the robustness of gradient shape are further enhanced if transverse diffusion effects are considered. We show that the positional error decreases as the tissue widens, but then saturates as the width reaches about ten cells. Our findings provide further evidence that noisy morphogen gradients are able to guide high-precision tissue patterning, and offer a reason why certain developing tissues may favor a specific dimension over another in the context of patterning precision.

## Results

### A 2D model for cellular tissue patterning

We approximate the cellular tissue with a regular rectangular 2D lattice of length *L*_*x*_ and width *L*_*y*_, such that patterning occurs along the *x*-axis. The lattice represents a 2D patch of cells of diameter *δ* each, which determines the domain length *L*_*x*_ = *N*_*x*_*δ* and width *L*_*y*_ = *N*_*y*_*δ*, given the number of cells *N*_*x*_ and *N*_*y*_ in *x* and *y* direction (Fig. 1B–D). We ignore variability in cell areas or diameters here, as physiological levels were previously shown to have negligible impact on gradient precision [18]. The patterning axis of length *L*_*x*_ is further divided into a morphogen source [−*L*_s_, 0] and a patterning domain [0, *L*_p_], such that *L*_*x*_ = *L*_s_ + *L*_p_. The steady-state reaction-diffusion equation from 1D (Eq. 3) is extended to 2D by writing

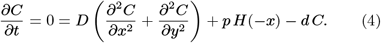

We solved Eq. 4 numerically with cell-to-cell stochasticity in the three kinetic patameters (see Methods for details), with two sets of boundary conditions (BCs): 1) zero-flux boundary conditions on all four sides of the domain, and 2) periodic BCs in *y* direction and zero-flux BCs in *x* direction. The latter is to mimic an infinitely wide domain and to minimize the finite size effects that result from the choice of specific BCs in *y*. While zero-flux boundaries along *y* may be the more relevant case for flat tissue geometries, we included periodic boundaries to demonstrate that our results are largely independent of the chosen BCs. A further motivation is that periodic BCs model tubular tissues that are patterned along their long axis.

### Gradients are more robust in wider tissues

The 2D solution can be viewed as an ensemble of gradients along each cell row, correlated by transverse diffusion. For a patterning domain with *N*_*y*_ cells in width, Eq. 1 can be fitted to each of the *N*_*y*_ cell rows. The variability in the gradient parameters λ and *C*_0_ is then quantified by the coefficient of variation CV_λ_ = *σ*_λ_*/μ*_λ_ and 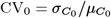 at a fixed position *y*, but over independent realizations of the 2D gradient. As the number of cells orthogonal to the patterning axis, *N*_*y*_, increases from 1 to 40 at fixed cell size, our stochastic gradient analysis shows that the variability of λ and *C*_0_ is inversely proportional to the square root of *N*_*y*_ (Fig. 2A,B,E,F):

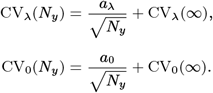

Specifically, under zero-flux boundary conditions and with a physiological amount of cell-to-cell kinetic variability (Methods), *a*_λ_ = 0.0294 ± 0.0004, CV_λ_(∞) = (9.96 ± 1.91) × 10^−4^, *a*_0_ = 0.2113 ± 0.0026 and CV_0_(∞) = 0.0028 ± 0.0012 as determined by curve-fitting (bounds are standard errors). This square-root relation can be explained by the Law of Large Numbers, which (among other things) states that the sampling variance of the mean of *N* identically distributed random variables is proportional to 1/*N*. In a 2D tissue, up to an offset accounting for the finite reach of transverse diffusion, the gradient variabilities can be understood to be sampling standard deviation of a correlated population of gradients, therefore scaling as 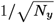. The offsets CV_λ_(∞) and CV_0_(∞) let the variability saturate as the domain width reaches infinity. The gradient variability parameters CV_λ_ and CV_0_ decrease significantly as the tissue width reaches about ten cells, before the gradient variability gradually flattens (Fig. 2A,B).

**Fig. 2:**
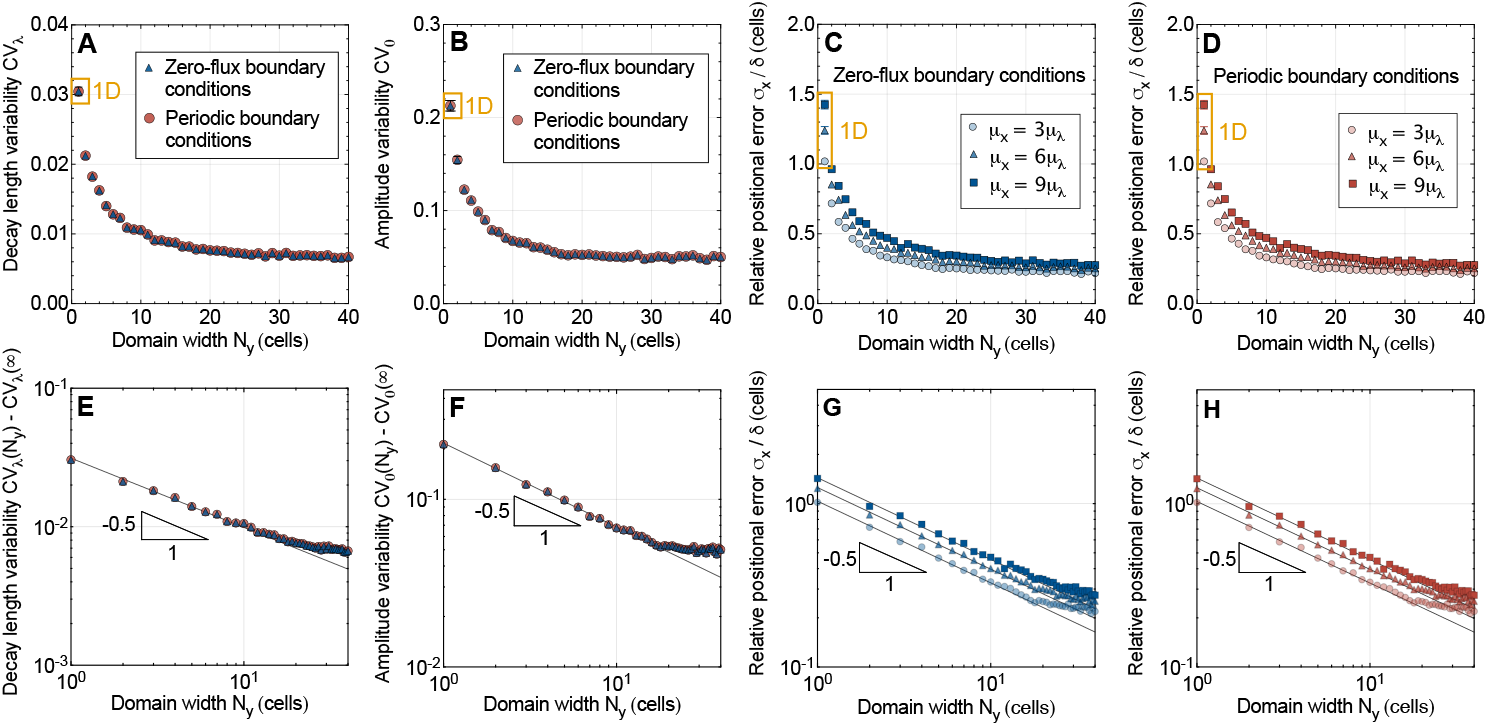
Impact of domain width on gradient variability and readout precision. Statistical evaluation of *n* = 1000 numerically generated, noisy morphogen gradients with cell-to-cell variability in the kinetic parameters *k* = *p, d, D* defined by a coefficient of variation CV_*k*_ = 0.3. The gradient variability and positional errors were calculated at a single cell-row at the middle of the domain (i.e., at *y* = *L*_*y*_/2). Error bars are SEM, but are mostly smaller than the symbols. Generated 2D gradients with a single cell row (*N*_*y*_ = 1, orange) are identical to 1D gradients within statistical errors. **A**,**B** Gradient variability, as measured by the coefficients of variation for the gradient decay length (CV_λ_) and amplitude (CV_0_) as a function of the domain width. Results for zero-flux (blue triangles) and periodic (red circles) BCs are nearly equal. **C**,**D** Positional error in units of cell diameters, measured at three different readout positions along the patterning axis (different symbols), as a function of the domain width, for zero-flux (C, blue) and periodic tissue boundaries (D, red). **E–H** Log-log plots of A–D reveal the square-root scaling of gradient variability and of the positional error with the domain width. Solid lines are fitted relationships of the form 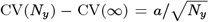 and 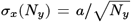. Deviation from the square-root scaling sets in as *L*_*y*_ exceeds a certain width (see also Supplemental Figs. S1 and S2).

Our findings echo with Bollenbach et al. [5] in that 2D tissues weaken the effect of deficient cells on morphogen concentration, thus lowering the concentration uncertainty. From this point of view, the wider the tissue, the larger the set of diffusion paths morphogens can choose from to bypass deficient cells, at the cost of a longer path to travel. Increasingly long detours gradually weaken this compensatory effect, such that the benefit of even wider tissues tends to zero. More generally, diffusion in *y* smooths out local fluctuations by allowing the noise experienced by one gradient to disperse among its neighbors, which is otherwise harder to achieve in 1D. Indeed, the gradient variability in wider tissues is considerably lower than that predicted in 1D (marked in orange in Fig. 2A–D) [17, 18].

### Patterning precision increases in wider tissues

Not only gradient variability is reduced, but also the spatial accuracy they convey to the patterned cells is enhanced in wider tissues (Fig. 2C,D,G,H). Similar to the scaling of gradient variability with respect to the domain width, the positional error is inversely proportional to the square root of the domain width at fixed cell size:

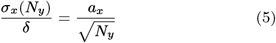

where *a*_*x*_ is a dimensionless factor discussed in detail in the following sections. Our data from *n* = 1000 two-dimensional morphogen gradients suggests that, unlike the gradient variabilities, the positional error does not possess a statistically significant intercept *σ*_*x*_(∞).

The further away from the morphogen source, the larger the positional error [5, 17, 18], an effect that we also observe here in 2D. Nevertheless, our gradient quantifications revealed that even for a readout position as far as *μ*_*x*_ = 9*μ*_λ_ (where *μ*_λ_ = 4*δ* is the average gradient decay length), the positional error is significantly reduced in wider domains. Eq. 5 holds independent from the position in the pattern (Fig. 2), and regardless of the boundary conditions.

In Eq. 5 and Fig. 2, we non-dimensionalized the positional error by the constant cell diameter *δ* to provide some intuition for the patterning precision relative to the size of the cells. Remarkably, extending the tissue width from one to two cells is sufficient to bring the positional error to subcellular levels even at the readout position furthest from the source, *μ*_*x*_ = 9*μ*_λ_, and in wider domains it can go considerably lower still. Specifically, for a nearly square tissue with 50 cells along the patterning axis and 40 cells in width, the positional error at *μ*_*x*_ = 3*μ*_λ_ is reduced to less than a fourth of a cell diameter (*σ*_*x*_/*δ* = 0.22). With a readout position far away from the source, i.e. at *μ*_*x*_ = 9*μ*_λ_, the positional error is still reduced to 28% of one cell diameter.

In the generation of these gradients, we fixed the cell diameter *δ* = 5 μm. Similar behavior is observed with larger and smaller cells, the only qualitative difference being that the deviation sets in at different number of cell rows *N*_*y*_, but at approximately equal domain widths *L*_*y*_ = *N*_*y*_*δ* (Supplemental Figs. S1 and S2). Moreover, we determined the gradient variability and the positional error between different tissues or embryos in the middle of the domain width, at *y* = *L*_*y*_/2, in Fig. 2. The results are unchanged if averaged over the entire width, as may be more relevant for robust development, except that the square-root scaling (Eq. 5) is found to hold even more robustly, over the entire range of tested tissue widths (Supplemental Fig. S3).

### Patterning precision increases with narrower cells

Cell shapes are not always isotropic. To test the effect of polarized cell shapes on patterning precision, we relaxed the previous assumption that cells have the same diameter *δ* in both tissue directions, and tested how the readout precision would be affected by the cell aspect ratio *r* = *δ*_*x*_/*δ*_*y*_, defined as the ratio between the cell diameters along the patterning axis and orthogonal to it (Fig. 3A). As the diameter along the patterning axis is fixed but cells widen orthogonal to it, we observe that the positional error increases (Fig. 3B,C). The cell aspect ratio affects the positional error, now a function of both cell diameters *δ*_*x*_ and *δ*_*y*_, as

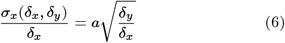

where *a* is a dimensionless proportionality constant. Narrower cells thus increase the precision of the positional information carried by the morphogen gradients, and this effect intensifies the narrower the cells become. The prefactor *a* encodes the scaling effect of other length scales in the patterning system, such as the mean readout position *μ*_*x*_, the average gradient decay length *μ*_λ_, and the source size *L*_s_. From 1D simulations, it is known that 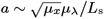 asymptotically [18], a relationship that we have verified to also hold in 2D analogously. Through the omitted prefactor, *a* will also depend on the type of 2D cell arrangement (here a rectangular array for simplicity), the morphogen sensing strategy employed by the cells [18] (here concentration averaging over the cell area), and possibly on other factors like the non-uniformity of cell sizes or the precise nature of the variability in the kinetic parameters.

**Fig. 3:**
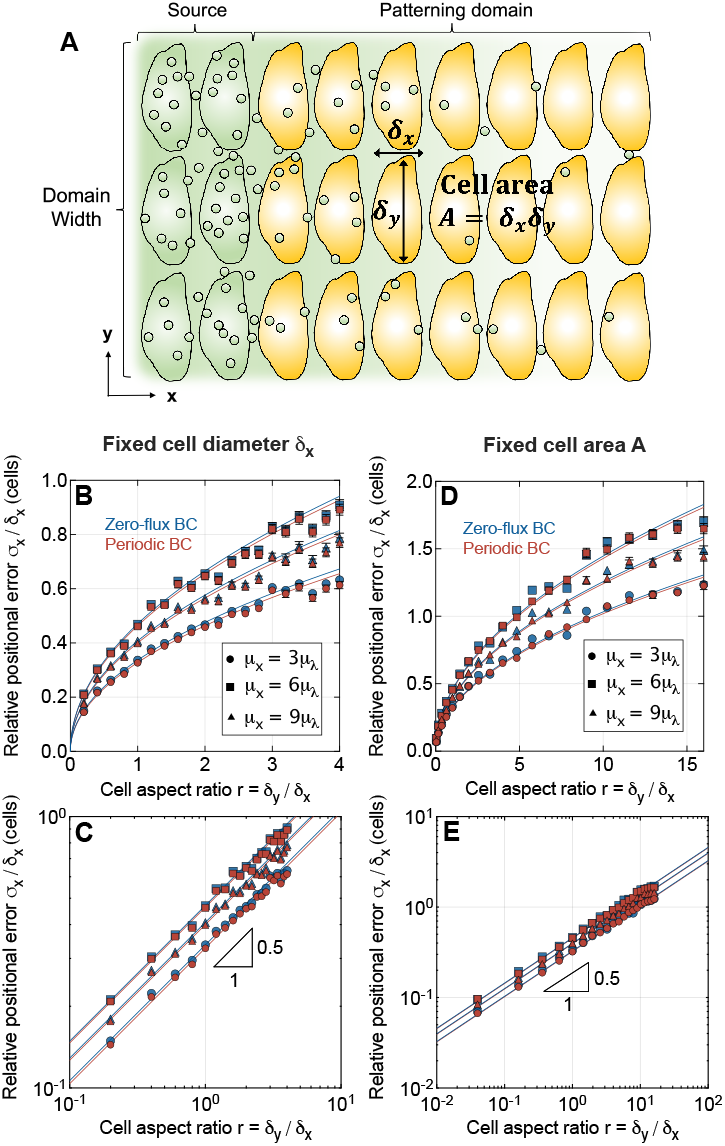
Effect of cell aspect ratio on patterning precision. **A** Schematic of a 2D tissue with anisotropic cell shapes. **B** When cells have a fixed diameter of *δ*_*x*_ along the patterning axis and a varying diameter *δ*_*y*_ in transverse direction, patterning precision decreases with wider cells (larger aspect ratio). **C** Log-log plot of the data shown in B, revealing a square-root scaling 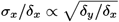 **D** Dependency of the positional error on cell aspect ratio for fixed cell area *A* = *δ*_*x*_*δ*_*y*_. **E** Log-log plot of D, revealing a square-root scaling as in C. Data points indicate mean ± SEM from *n* = 1000 independent gradient realizations. Different symbols represent different readout positions along the patterned axis. Solid lines are fitted square-root relationships.

As Eq. 6 shows, *relative* to the cell diameter in patterning direction, it is the cell *shape* that determines the patterning precision, through its aspect ratio. In *absolute* terms, on the other hand, it is the cell *area* instead: Eq. 6 can be rearranged to

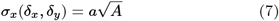

where *A* ∝ *δ*_*x*_*δ*_*y*_ is the cell area. Fixing the aspect ratio but changing the cell area alters the *absolute* positional error, but not the *relative* one. Keeping the cell area constant while varying the aspect ratio, on the other hand, alters the *relative* positional error (Fig. 3D,E), but not the *absolute* one.

In summary, not only cell size, but also cell shape affect the patterning precision in a 2D tissue, but it is a matter of perspective which of the two does. The absolute positional error carried by morphogen gradients in a two-dimensional patterning scenario with kinetic cell-to-cell variability is lower if cells are narrower not only along the direction of the gradient, but also if they are narrower in the transverse direction (Eq. 7). This can intuitively be understood from the smoothing effect that transverse diffusion has on morphogen levels.

### Fast transverse diffusion increases patterning precision in wide tissues

Given the buffering effect that transverse diffusion has on morphogen gradients as shown above, it is natural to ask whether a variability reduction can be achieved with anisotropy in the morphogen diffusivity. While the relevance of anisotropic diffusion for gradient-based patterning in specific developing tissues remains to be studied experimentally, we will provide a theoretical analysis. We restrict our analysis to the simplest case with a diagonal diffusion matrix aligned with the *x* and *y*-axes of the tissue. This introduces a single new parameter, the degree of diffusion orthotropy, ∝ = *D*_*y*_ /*D*_*x*_ (see Eq. 8 in Methods). Intuitively, the noise buffering effect is expected to be boosted by faster transverse diffusion. Indeed, varying *α* in simulations, we observed a reduction both in the gradient variability and in the positional error as the transverse diffusivity becomes stronger (Fig. S4). Naturally, this effect is observed only for *N*_*y*_ > 1 in our model. Asymptotically toward wide tissues (large *N*_*y*_), we found that the variabilities and positional error are reduced proportionally to 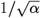 *α* as expected from the 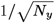 scaling found in Fig. 2. In combination, we thus have

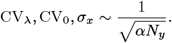

Altogether, we arrive at the overall positional error of noisy exponential morphogen gradients in 2D epithelia of sufficient width,

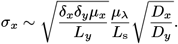

Thus, our theoretical model predicts that patterning precision does not only increase with smaller cells, but also with faster transverse diffusion in wide enough tissues. Notice that ultimately, the cell width *δ*_*y*_ affects the positional error through the number of cells along the tissue width, *N*_*y*_ = *L*_*y*_ /*δ*_*y*_.

### Patterning precision in the *Drosophila* wing disc

In the *Drosophila* wing disc (Fig. 4A), the Dpp gradient defines the positions of the longitudinal veins. Strikingly, Dpp readout positions remain at the same relative location in the domain despite substantial tissue growth, a phenomenon that is referred to as pattern scaling [22, 30]. The position of the anterior Spalt domain boundary, which places the L2 vein, remains at 40–45% of the anterior domain length [31, 5, 30]. Scaling of the readout with the length of the patterning domain is achieved by the parallel increase in the gradient amplitude and length [22, 32]. We previously showed that the positional error remains largely constant in spite of the growing absolute readout distance from the source and the expanding gradient length, because the parallel increase in the source width and decrease in the average apical cell areas [33–35, 22, 36] compensate according to 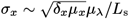 [18]. While the predicted positional error at *μ*_*x*_*/L*_p_ = 0.4 remains below about 4 μm when considering a 1D domain [18] (Fig. 4B, green), this still corresponds to more than two cell diameters. With a domain that is *N*_*y*_ = 10 cells in width—a number easily surpassed in the wing pouch—we find a positional error well below a single cell diameter throughout development (Fig. 4B, orange) with isotropic 2D diffusion, a more-than-3-fold reduction compared to the 1D perspective. These observations suggest that 2D effects are key to ensuring high patterning precision in tissue development.

**Fig. 4:**
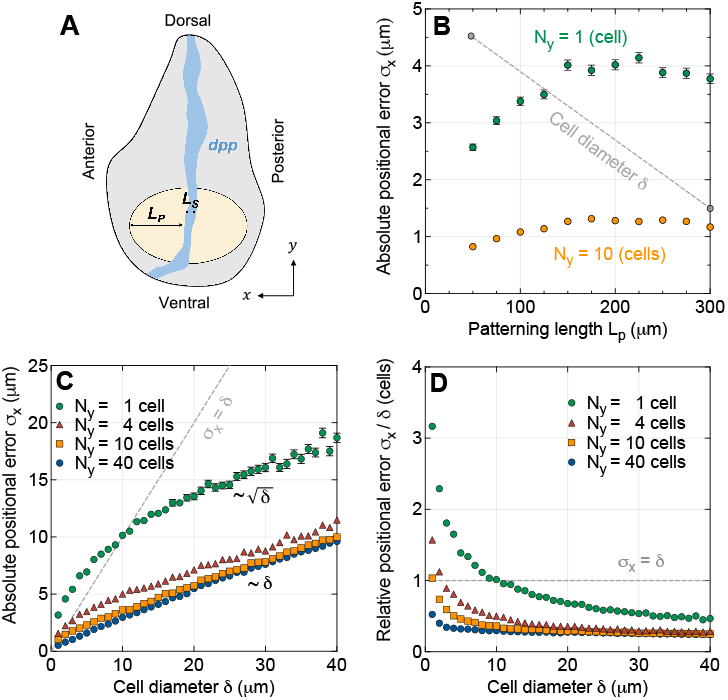
Effect of tissue width on patterning precision in developing tissues. **A** Schematic illustration of the *Drosophila* wing disc (gray) with its Dpp source along the anterior-posterior boundary (blue). The tissue width orthogonal to the Dpp gradient is in dorsal-ventral direction. The wing pouch is shown in beige. **B** Evolution of the predicted positional error of the Dpp gradient as the wing pouch widens over time, at *μ*_*x*_ = 0.4*L*_p_ for *μ*_λ_ = 0.11*L*_p_, *L*_s_ = 0.16*L*_p_, and *δ*_*x*_ = 5.1 μm − 0.012*L*_p_ [18]. When the tissue width is accounted for by 10 cell rows (*N*_*y*_ = 10, orange), the positional error is substantially lower than with only one cell row (*N*_*y*_ = 1, green), and remains well below once cell diameter (gray dashed line). **C** Comparison of absolute positional errors at *μ*_*x*_ = 9*μ*_λ_ for varying tissue widths, as a function of cell size, for isotropic cell shapes and diffusion. **D** Comparison of positional errors as in C, relative to the cell diameter. Data points indicate mean ± SEM from *n* = 1000 independent gradients.

### Patterning with subcellular precision is possible in wide tissues irrespective of cell size

The absolute positional error grows in proportion to the square root of the cell area (Eq. 7). In epithelia that are patterned by morphogen gradients, cells have indeed been found to possess smaller apical areas than in epithelia that are not [18]. This raises the question whether morphogen gradient-based patterning can still be sufficiently precise in tissues with larger or less densely packed cells, such as mesenchymes. The reported cell densities of 11–28 cells/1000 μm^2^ [37] imply a cell diameter of 6–10 μm (including extracellular matrix) in the chick forelimb bud at Hamburger–Hamilton (HH) stages 18–25. Images of the chick forelimb bud at HH 21 [38] suggest a similar cell diameter of about 8–10 μm. Compared to the 5 μm of neuro-epithalial cells [6], this would entail a 1.2–2-fold reduction in absolute patterning precision. In units of cell diameters, however, the precision remains unaltered (Eq. 6), ensuring that the right cell numbers choose their fate according to plan also when they are large.

The absolute positional error grows with increasing cell diameter, i.e., 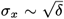 in 1D [18]. Given the square root relationship, the positional error will exceed a cell diameter for small cells, but for sufficiently large cells, the opposite is the case (Fig. 4C). Patterning with subcellular precision is possible beyond a cell diameter of about 10 μm in a quasi-one-dimensional tissue (Fig. 4D). In 2D, this relationship transitions to linear, i.e., *σ*_*x*_ ∝ *δ* for *δ* = *δ*_*x*_ = *δ*_*y*_ (Eq. 7, Fig. 4C). The crossover point *σ*_*x*_ = *δ* is attained at a smaller cell diameter, the wider the tissue. Already with *N*_*y*_ = 4 cells, we observe a subcellular positional error at cell diameters of about 2.5 μm or greater (Fig. 4D). In developing tissues that are more than ten cells wide—which they commonly are—subcellular patterning resolution is obtained even with cells as small as *δ* = 1 μm or greater. The precision-enhancing buffering effect in 2D makes high-precision patterning possible in tissues covering a wide range of cell sizes.

## Discussion

Molecular noise has long been believed to cause large variability during embryonic development. This has started to change recently, and our present findings underpin this trend. By statistically analyzing the effect of molecular noise on morphogen concentration gradients, we show that for realistic morpho-kinetic noise levels, positional information can be expected to be conveyed to cells in two-dimensional tissues with sub-cellular resolution. While our study builds on recently developed modeling ideas [17, 18], a new perspective was required to arrive at this result: Tissue patterning typically occurs in at least two dimensions, and morphogens can diffuse also orthogonal to the patterning axis. Not only does this allow them to bypass malfunctioning cells [5], transverse diffusion also substantially smoothens out local morphogen fluctuations in the entire tissue, leading to a considerable drop in gradient variability and in the positional error.

We found that patterning precision scales with the square root of the number of cells along the tissue width—an intuitive relationship with theoretical roots in statistics. Arriving at this result required extending the previous 1D model for patterning to two dimensions. A striking consequence of this relationship is that even a few cells across are sufficient to allow morphogen-sensing cells to differentiate according to their location in the pattern with a positional accuracy below a single cell diameter. Compared to 1D patterning scenarios, 2D morphogen transport allows for a considerable precision enhancement up to a tissue width of roughly ten cells, before saturation sets in. The positional error in 2D tissues differs from the 1D expression [18] in that it contains an additional scale encoding the tissue width in units of cells, and the degree of anisotropy in the morphogen diffusivity:

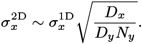

In wide enough tissues, the precision gain can be as big as 3- to 4-fold even with isotropic diffusion, which further reinforces recent results suggesting that morphogen gradients in the vertebrate neural tube carry more precise spatial information for patterning than previously thought [17]. While a 1D noisy gradient model just explained the observed accuracy of the central readout boundaries in neural tube patterning, 2D diffusion allows that accuracy to be surpassed even, leaving wiggle room for additional sources of noise not accounted for by our stochastic gradient model. The precision-enhancing 2D effects observed here are not limited to a close vicinity of the morphogen source, but they apply anywhere in the tissue, also in regions distal from the source. Our data suggests that high-precision tissue patterning with morphogen gradients is possible in spite of molecular and biological variability also far away from the origin.

Our work also reveals how morphogen gradient variability depends on the cell area and aspect ratio in epithelia. The absolute positional error is proportional to the cell area. Therefore, the patterning precision increases when cells are narrower in both tissue directions. However, we found that this dependency applies only to the *absolute* error *σ*_*x*_. For robust and precise tissue development on the cellular level, it may be relevant to let cells differentiate in the right numbers, and so the positional error in units of the cell size in patterning direction, *σ*_*x*_/*δ*_*x*_, may be the relevant quantity instead. Strikingly, we found this *relative* error to depend on the cell aspect ratio instead of the cell area. The wider the cells are compared to their length along the patterning axis, the larger the positional error in terms of numbers of cells. This may equip Nature with a tunable degree of freedom that can be tailored to a required level of developmental precision—a hypothesis that could be amenable to experimental testing, and which could be relevant for gradient-based tissue engineering.

Our findings also hint at a developmental advantage for widening tissues, as long as their cells do not widen with it. What brings down the positional error is the number of cells across the tissue, *N*_*y*_, not the width itself, *L*_*y*_, if noise in the gradients indeed arises from inter-cellular kinetic variability. These results put our previous analysis on the effect of cell size on patterning precision [18] into a broader context. In the *Drosophila* eye disc, the initial size at 40 h after hatching is about [*L*_*x*_, *L*_*y*_] = [50, 100] μm [39]. With an apical area of 7 μm^2^ and smaller [34], 100 μm corresponds to at least *N*_*y*_ = 60 cells in width. We now find that for such tissues, the positional error is reduced by about 4-fold compared to the traditional 1D perspective. For larger number of cells, the precision-enhancing effect, however, plateaus. Francis Crick started his seminal paper on the role of diffusion in gradient patterning by quoting Lewis Wolpert, “It has been a great surprise and of considerable importance to find that most embryonic fields seem to involve distances of less than 100 cells, and often less than 50.” [20]. In light of our work, such tissue widths appear sufficient for precise patterning.

Although we linked the implications of 2D diffusion for patterning precision to two specific epithelial tissues—the vertebrate neuroepithelium and the fly wing disc—the main results presented here are system-agnostic, and might underlie developmental precision across species more generally. As with all theoretical and modeling work, our predictions from a statistical analysis of synthetically generated noisy morphogen gradients requires experimental validation, which we hope to motivate herewith. An open question is the impact of complex tissue architecture, such as that of pseudostratified epithelia [40], on patterning precision [27]. In light of our study, it appears plausible that the densely packed cells with stratified nuclei can help buffer morphogen level fluctuations similar to the stratified cell rows in our 2D cell arrays here, at least for morphogens that are sensed along the lateral sides of cells, such as Bone Morphogenetic Protein (BMP) in the neural tube [41]. Our results would then suggest that the positional error cells experience from reading out the BMP gradient is reduced by the strong pseudostratification. Generally, one may ask if (and to which degree) patterning precision can be further enhanced in three-dimensional tissues, a question that we leave for future research. While our model is straightforward to generalize to 3D, previous results by Bollenbach et al. [5] hint at an affirmative answer.

Finally, a few potential limitations of our model deserve being mentioned. While our statistical approach to generate noisy morphogen gradients is among the ones with the most detailed representation of molecular noise in the literature, there can be a range of other sources of biological variability involved in tissue patterning processes that were not represented here. Noise in the molecular readout mechanism, in regulatory interactions (either during the formation of the morphogen gradient or down-stream), or in the fate decision process, are some of them. We restricted our analysis of spatial precision to the pure gradients here. Moreover, our analysis is based on a representation of the morphogen concentration as a continuous function *C*(*x, y*), ignoring the discrete, particulate nature that may become relevant at low morphogen numbers. Variability in the cell areas, on the other hand, would not be expected to alter our findings [18], although it may be of interest to extend the present model from a rectangular cell grid to more realistic tissue geometries with irregular cell arrangements.

## Methods

### Generation of 2D morphogen gradients from molecular noise

To study the inter-embryonic variability and intra-embryonic noisiness of morphogen gradients in a quantitative way, we generated large numbers of statistically independent gradients numerically. We discretized the 2D tissue domain [− *L*_s_, *L*_p_] × [0, *L*_*y*_] into a rectangular array of biological cells (Fig. 1B). The standard setup consisted of five columns of cells making up the morphogen source domain (*x <* 0), and 50 columns in the patterning domain adjacent to it, where the morphogen gradient forms (*x* ≥ 0). To observe the effect of tissue width, the number of cell rows was varied in the range 1– For each cell at position *i, j* in the tissue, we drew random morpho-kinetic parameters *p*_*ij*_, *d*_*ij*_, *D*_*ij*_ from log-normal distributions analogous to previous 1D studies [17–19], such that the noisy exponential morphogen gradients *C*(*x, y*) emerged as approximate solutions of the 2D steady-state reaction-diffusion problem

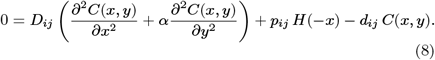

The parameter *α* represents the degree of orthotropy in morphogen diffusion, which we set to 1 (isotropic diffusion) except where the effect of polarized morphogen transport was analyzed explicitly. Each of the kinetic parameter distributions was defined by a mean value *μ*_*k*_ and a coefficient of variation CV_*k*_ = *σ*_*k*_/*μ*_*k*_ for *k* = *p, d, D*. Physiological molecular noise levels were identified previously [17], and we used the same value CV_*k*_ = 0.3 here. We then solved the PDE (Eq. 8) with the finite difference method, using a five-point central difference stencil, in Matlab 2021b. Each biological cell used 3 ×3 finite difference grid points, which offered a good compromise between computational speed and numerical accuracy.

For mass conservation, we imposed continuity of the morphogen concentration and flux fields everywhere in the 2D domain. Note that in a strict mathematical sense, this implies that Eq. 8 can only be approximated, but not solved exactly, in dimensions higher than one and with uncorrelated random diffusivities *D*_*ij*_. This is akin to the situation in reality, in which the diffusivity field will not be a perfect step function as used here, with discontinuities at the cell boundaries. Within spatial accuracy of the finite difference discretization, the numerical approximation of Eq. 8 automatically smoothens the parameter fields to enable a mass-conserving solution.

Zero-flux boundary conditions were imposed in *x* direction, and zero-flux or periodic boundaries were assumed in the width direction (*y*) as indicated in the figures. A relative error tolerance of 10^*−*10^ was used to terminate the iterative solver. Repeating this process with new random kinetic parameters for all cells yielded *n* = 1000 independent realizations of 2D morphogen concentration profiles (as shown exemplarily in Fig. 1C,D), which we then evaluated statistically.

### Calculation of gradient variability

To extract the amount of variability in the gradient decay length and amplitude, we sliced the 2D morphogen concentration profiles *C*(*x, y*) along rows of cells *j* in the direction of patterning. These 1D slices *C*_*j*_ (*x*) of the gradients are spatially correlated due to transverse diffusion, and were approximately exponential along *x*, with a slight bend at the distal tissue boundary due to the imposed boundary conditions. Along the central cell row *j* = *N*_*y*_*/*2 (or along all of them for Supplemental Fig. S3), we therefore fitted the deterministic homogeneous solution [17]

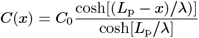

to the noisy concentration profiles in the patterning domain, after logarimizing them first. Repeating this process for all *n* 2D gradients yielded a set of statistically independent gradient decay lengths {λ_1_,..., λ_*n*_} and amplitudes {*C*_0,1_,..., *C*_0,*n*_ }, each representing one embryo or tissue, from which we computed the coefficients of variation, CV_λ_ and CV_0_. Standard errors were determined using bootstrapping.

Note that the coefficients of variation CV_*p,d,D*_ in the kinetic parameters quantify inter-cellular variability that governs the gradient noisiness *within* embryos, but also determine the variability thus emerging *between* them, as measured by CV_λ_ and CV_0_.

### Statistical evaluation of readout positions and patterning precision

We used spatial averaging over the cell surfaces to determine the morphogen concentration each cell is exposed to—a choice that does not affect the positional information conveyed to the cells significantly [18]. With the resulting concentration profiles in the form of step functions, we determined the positional error according to its definition, as the standard deviation of the locations where the gradients attain a threshold value (Eq. 2). Since 2D gradients are not always monotonic, there can be multiple such locations along a single gradient. In such cases, we used their mean location to define the readout position *x*_*θ*_. The average readout position was then determined by averaging over all gradient realizations, *μ*_*x*_ = mean[*x*_*θ*_]. At three different such (average) read-out positions *μ*_*x*_ = 3*μ*_λ_, 6*μ*_λ_, 9*μ*_λ_, we evaluated their standard deviations *σ*_*x*_ = stddev[*x*_*θ*_] over the *n* = 1000 noisy gradients. Standard errors were then again calculated using bootstrapping.

## Code Availability

The source code is released under the 3-clause BSD license. It is available as a public git repository at https://git.bsse.ethz.ch/iber/Publications/2023_long_2d_precision.

## Acknowledgements

This work was partially funded by the Swiss National Science Foundation under Sinergia grant CRSII5 170930.

## Author Contributions

Conceptualization, R.V. and D.I.; Software & Simulations, Y.L. and R.V.; Formal Analysis, R.V.; Supervision, R.V. and D.I.; Visualization, Y.L.; Writing, R.V., D.I., and YL.; Funding Ac-quisition, D.I.

---

**Figure S1:**
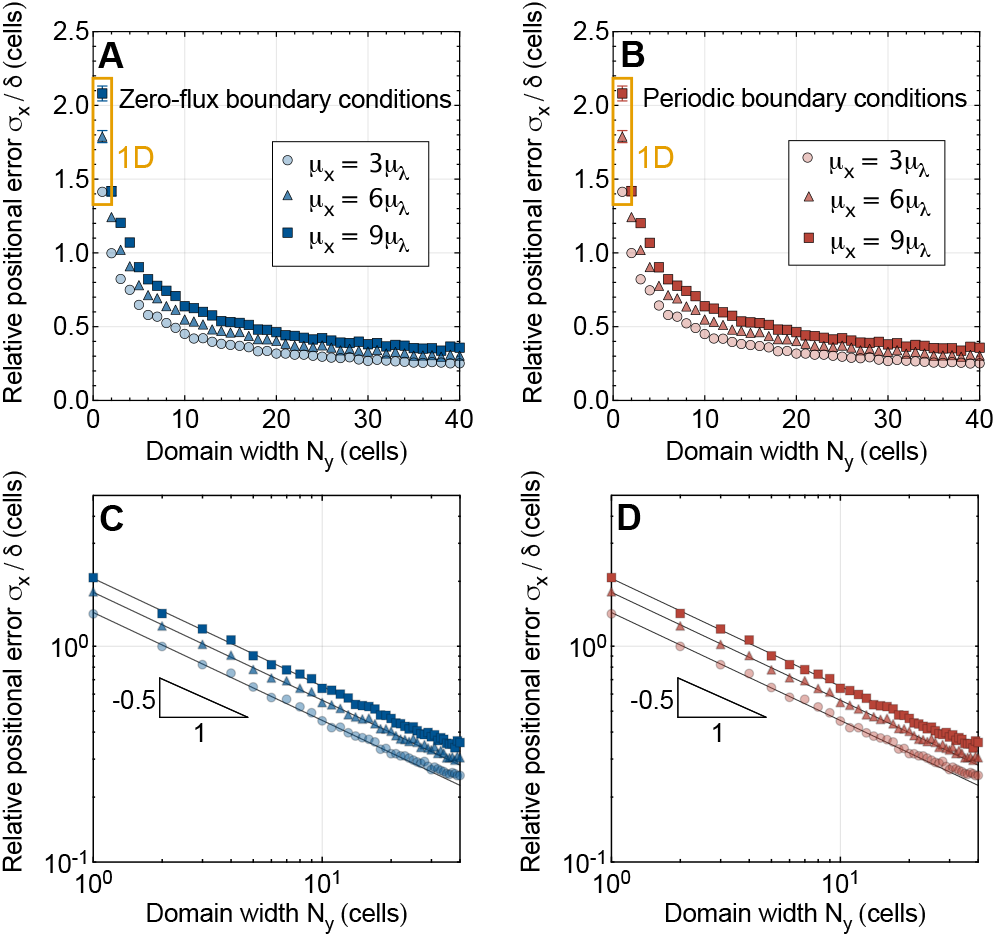
Impact of domain width on gradient readout precision for small cell diameters. Simulations analogous to those shown in Fig. 2C,D,G,H, except that the cell diameter is halved to *δ* = 2.5 μm. See caption of Fig. 2 for details.

**Figure S2:**
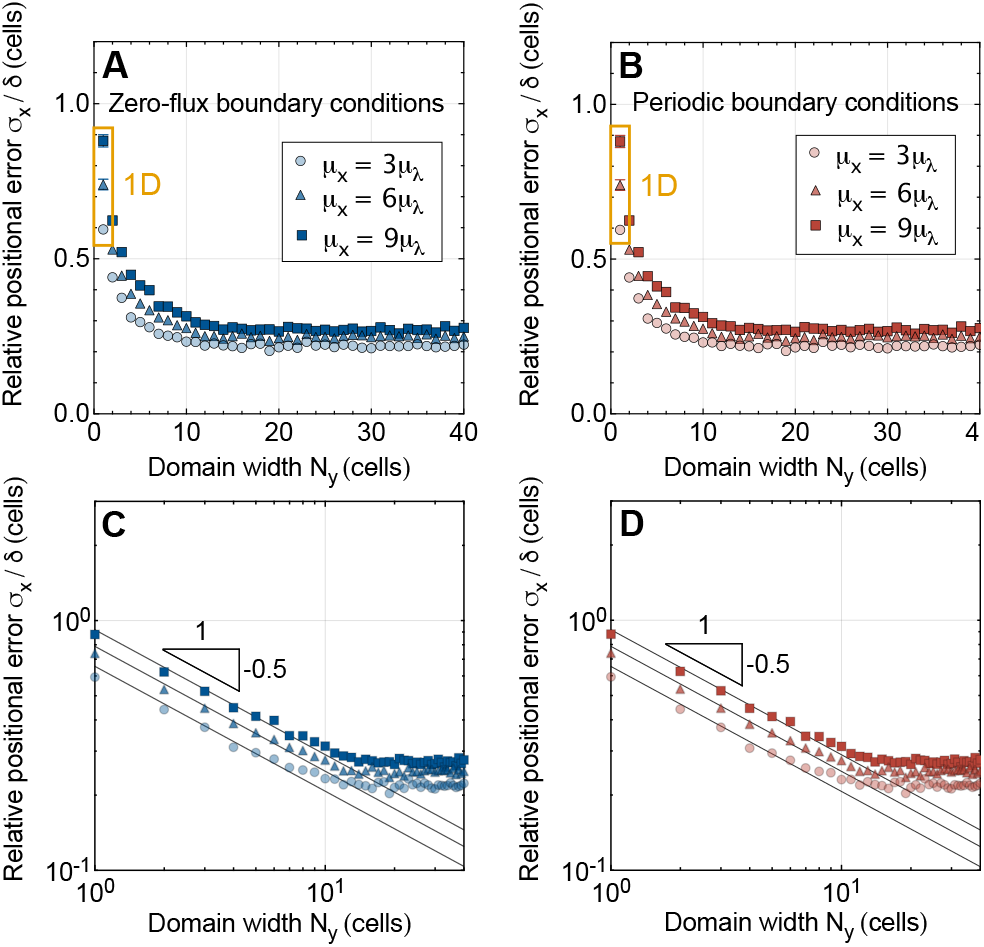
Impact of domain width on gradient readout precision for large cell diameters. Simulations analogous to those shown in Fig. 2C,D,G,H, except that the cell diameter is increased to *δ* = 12.5 μm. See caption of Fig. 2 for details.

**Figure S3:**
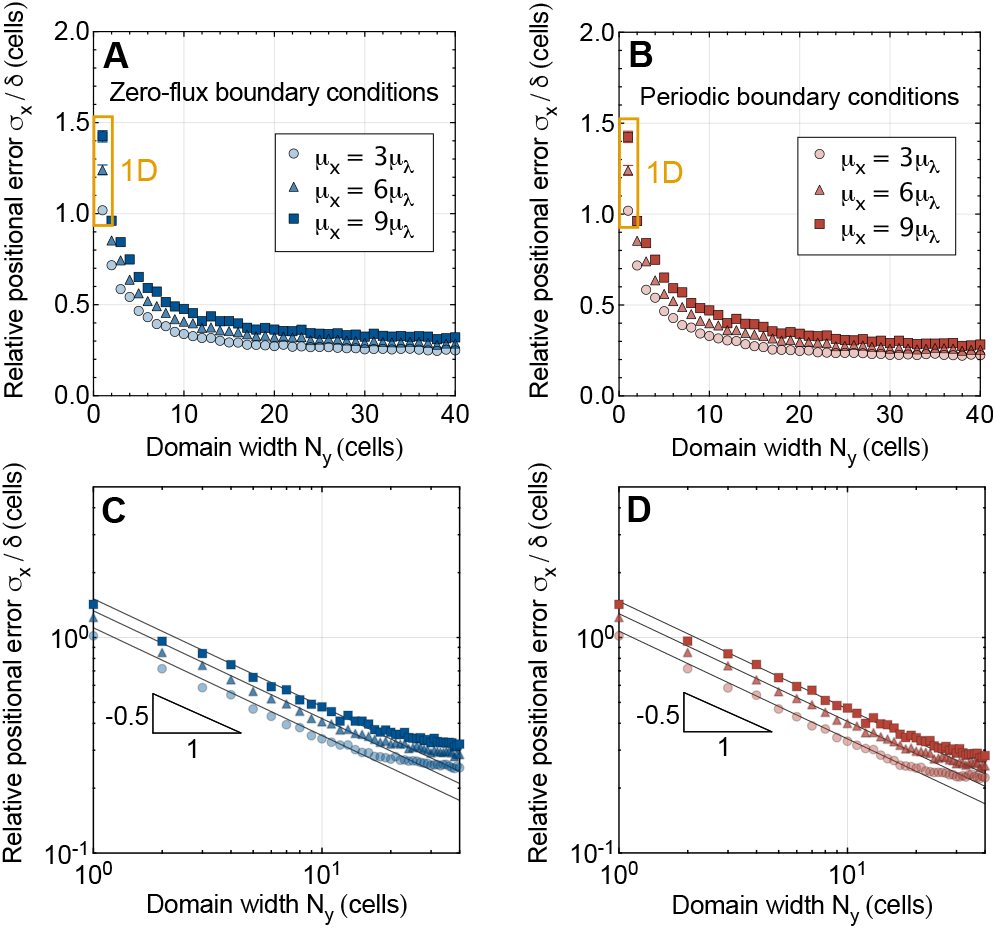
Impact of domain width on gradient readout precision averaged across the domain width. Simulations analogous to those shown in Fig. 2C,D,G,H, except that the positional error averaged over all cell rows is plotted. See caption of Fig. 2 for details.

**Figure S4:**
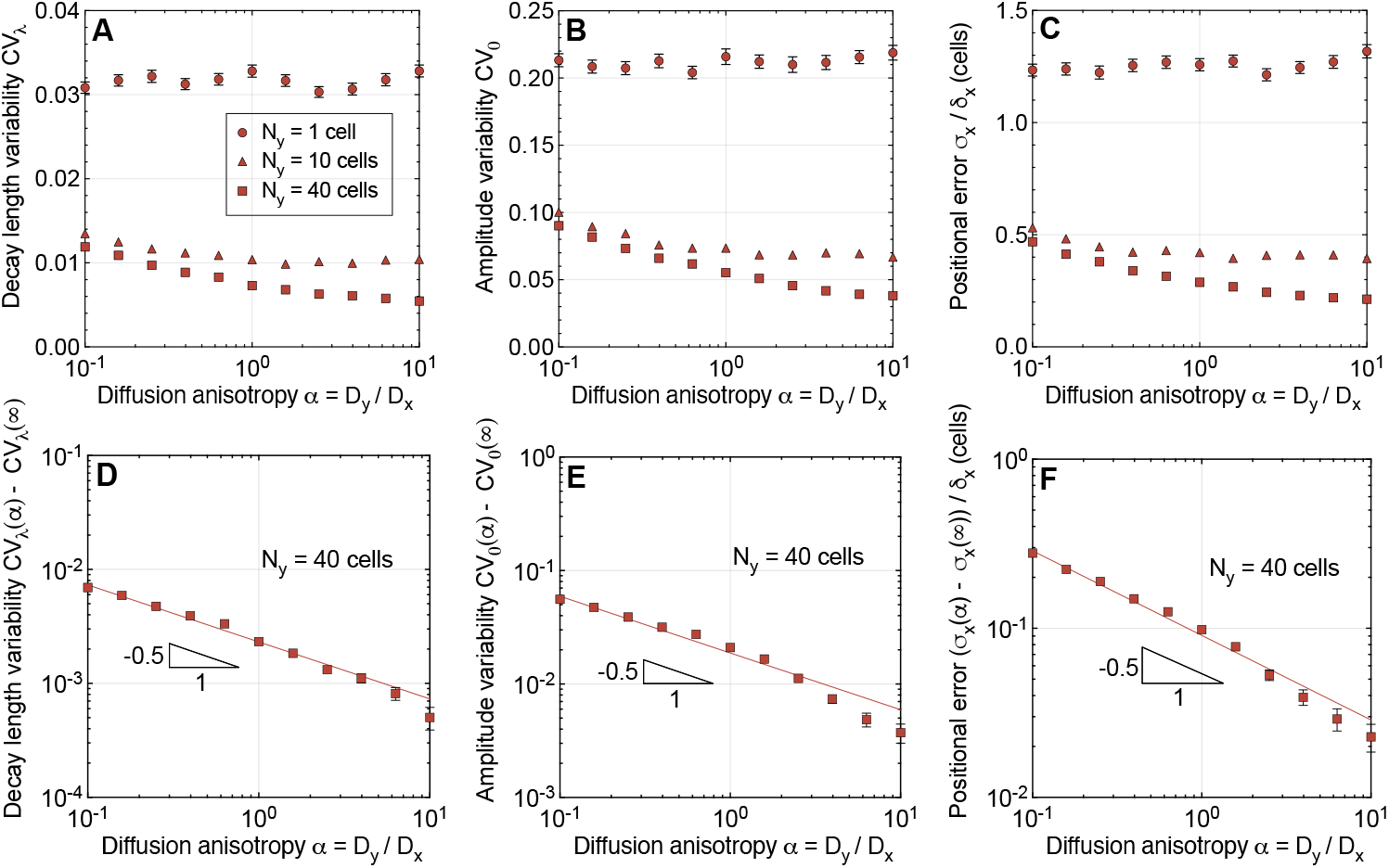
Effect of anisotropic morphogen transport on patterning precision. **A**,**B** Gradient variability as a function of the degree of orthotropy in morphogen transport, *α* = *D*_*y*_/*D*_*x*_. **C** Positional error at *μ*_*x*_ = 6*μ*_λ_ as a function of *α*. **D–F** Log-log plots of A–C at a tissue width of *N*_*y*_ = 40 cells, revealing square-root scaling when *N*_*y*_ is large enough. Data points indicate mean ± SEM from *n* = 1000 independent 2D gradients obtained on a tissue of square cells (*δ*_*x*_ = *δ*_*y*_).

